# Vessel Spatial Analysis (VeSpA): a tool for whole slide image segmentation, morphometry, and QuPath extension

**DOI:** 10.64898/2026.06.15.732366

**Authors:** Giulia Grion, Rash Hussain, Filippo Emanuele Colella, Kirollos Roufail, Silvia Uccella, Roberta Frapolli, Cristina Matteo, Ömer Mintemur, Francesca Pennati, Salvatore Lorenzo Renne

## Abstract

Quantifying vascular architecture in histological whole slide images is needed to study tissue organisation, tumour microenvironment biology, and disease-associated vascular remodelling. However, vessel analysis in routine immunohistochemistry remains challenging. Available workflows are often manual, require programming expertise, or lack direct integration with digital pathology platforms. We developed VeSpA (Vessel Spatial Analysis), an open-source pipeline and QuPath extension for automated vessel segmentation and morphometric quantification in CD31-stained whole slide images. VeSpA combines configurable signal extraction, using CMYK Yellow channel extraction by default and optional DAB stain deconvolution for H-DAB images, with automatic or percentile-based thresholding, morphological refinement, contour filtering, and lumen filling to generate vessel masks from standard DAB-stained sections. The QuPath extension includes a graphical interface for selecting annotations, TMA cores, or whole images, configuring segmentation parameters, running the Python backend, and importing vessel objects directly into the QuPath hierarchy. For each detected vessel, VeSpA extracts area, major axis length, minor axis length, eccentricity, centroid, and orientation, while also appending summary measurements to parent annotations and TMA cores. Validation against independent pathologist annotations showed that VeSpA achieved segmentation performance close to inter-rater agreement and outperformed yellow channel prompt-based SAM and zero-shot YOLOv8-seg on overlap-based metrics in the tested dataset. VeSpA integrates vessel segmentation, morphometric feature extraction, and QuPath-based visualisation into a single reproducible workflow for vascular quantification in computational pathology and spatial analysis of histological tissue architecture.

## 1 Introduction

The vasculature is a central component of tissue organisation and disease biology, supporting oxygen and nutrient delivery, immune cell trafficking, waste removal, and stromal communication [1, 2]. In cancer and other pathological conditions, vascular remodelling contributes to tumour progression, immune regulation, hypoxia, treatment resistance, inflammation, and fibrosis [3–8]. For this reason, quantitative assessment of vascular architecture in histological tissue sections can provide information beyond qualitative microscopic review.

Digital pathology has made it possible to analyse whole slide images (WSI) at scale, and platforms such as QuPath are now widely used for annotation, visualisation, cell detection, and biomarker quantification [9–13]. However, vascular analysis in routine immunohistochemistry (IHC) remains less standardised. Many workflows still rely on manual assessment, microvessel density estimation, or bulk stained-area measurement, while tools that segment individual vessels and extract vessel-level morphometric features directly within a digital pathology platform remain limited [14–16].

Existing vascular image analysis tools have advanced vessel and network quantification in several imaging contexts, including fluorescence microscopy, three-dimensional (3D) models, and specialised imaging modalities [15–22]. However, many of these approaches are not directly designed for standard two-dimensional (2D) formalin-fixed paraffin-embedded (FFPE) WSIs stained with chromogenic endothelial markers such as CD31. Deep learning methods can achieve strong performance when trained on appropriate annotations, but they require curated training datasets, computational resources, and domain adaptation that may not be available in many pathology research settings [23–27].

To address this gap, we developed Vessel Spatial Analysis (VeSpA), an open-source Python pipeline and QuPath extension for automated vessel segmentation and morphometric quantification in CD31-stained WSIs [12]. VeSpA was designed to operate on routine diaminobenzidine (DAB)-stained FFPE sections without requiring model training, GPU infrastructure, or command-line expertise. The tool segments vessel objects, extracts morphometric descriptors, imports results into the QuPath hierarchy, and supports validation against pathologist annotations and benchmarking against general-purpose segmentation models.

## 2 Materials and Methods

### 2.1 Image acquisition and histological processing

Histological FFPE tissue sections were stained by IHC using an anti-CD31 antibody to label vascular endothelial cells. Stained slides were digitised at high resolution using a whole slide scanner to generate TIFF image files. The pipeline can process any image format natively supported by QuPath, including pyramid WSI formats such as .svs,

.ndpi, .tiff, and .czi, as well as standard raster formats such as PNG and JPEG.

### 2.2 Software environment

The VeSpA pipeline is implemented in Python (v3.12.1), with a QuPath extension developed in Java for interactive use within the QuPath environment [12]. Key Python dependencies include OpenCV (v4.9.0.80), scikit-image (v0.22.0), NumPy (v1.26.3), pandas (v2.2.0), and Pillow (v10.2.0). All analyses were run on macOS 15.7.4 (24G517) with a 1.1 GHz Intel Core i3 dual-core processor. The Python code is available through a dedicated GitHub repository and is accompanied by online documentation. The QuPath extension is distributed separately and can be installed directly into the QuPath environment.

### 2.3 Python vessel segmentation pipeline

The core VeSpA segmentation workflow is implemented as a Python pipeline that takes CD31-DAB-stained histological images as input and generates lumen-filled vessel masks, visualisation overlays, and vessel-level morphometric measurements. The pipeline consists of two sequential modules: vessel segmentation and morphometric feature extraction. Vessel segmentation is performed in six steps: signal extraction, thresholding, morphological refinement, contour detection and filtering, lumen detection and filling, and mask generation.

#### Signal extraction

Images were loaded in BGR format using OpenCV. By default, VeSpA computes the Yellow (Y) channel of the CMYK colour model pixel-wise from the BGR image as follows:

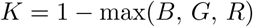

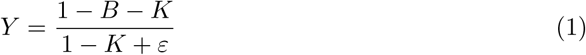

where *B* is the blue channel value, *K* is the key component, and *ε* = 10*^−^*^10^ is a small constant added to avoid division by zero. The resulting Y channel is scaled to the range [0, 255] and stored as an 8-bit unsigned integer array.

As an alternative preprocessing mode, VeSpA can also apply Haematoxylin (H)-DAB stain deconvolution and isolate the DAB channel as a stain-separated optical density signal. This DAB signal is normalised to an 8-bit grayscale image so that stronger DAB-positive regions correspond to higher foreground intensity prior to thresholding. Both signal-extraction modes feed into the same downstream thresholding, morphological refinement, contour filtering, lumen filling, and measurement workflow.

#### Thresholding

By default, Otsu’s thresholding algorithm was applied to the selected extracted signal, producing a binary mask in which vessel pixels are represented as white (foreground, value 255) and background pixels as black (value 0) [28]. As an alternative, a user-specified percentile threshold can be applied, where the threshold value is set to the *n*-th percentile of the extracted signal intensity distribution.

#### Morphological refinement

The binary mask was processed with two sequential morphological operations. First, dilation was applied using an elliptical structuring element of 21 *×* 21 pixels for one iteration. Second, erosion was applied using an elliptical structuring element of 3 *×* 3 pixels for two iterations.

#### Contour detection and filtering

External contours were extracted from the morphologically refined mask using cv2.findContours with the RETR EXTERNAL retrieval mode. Contours enclosing an area below a minimum threshold (500 px^2^ by default) were discarded; only qualifying contours were filled onto the output mask.

#### Lumen detection and filling

Vessel lumens were detected and filled in four sub-steps. First, fragmented vessel walls were repaired via morphological closing using an elliptical kernel (28 *×* 28 px, 2 iterations). Second, a flood-fill operation was initiated from the image border to identify true background pixels; candidate lumen regions were defined as pixels that were neither background nor vessel wall. Third, candidate regions were filtered by area (200–80,000 px^2^), eccentricity (*≤*0.97), and circularity (*≥*0.20), computed as 4*πA/P* ^2^. Fourth, validated lumen masks were expanded by a small dilation (5 *×* 5 px kernel, 3 iterations) and merged onto the original wall mask, ensuring precise contour preservation.

#### Mask generation and contour visualisation

The lumen-filled vessel mask constitutes the final segmentation output. A visualisation overlay was generated by blending the original histological image (weight 0.7) with a green-channel representation of the filled mask (weight 0.3) using cv2.addWeighted. Both the filled binary mask and the overlay were saved as PNG files in a dedicated subdirectory for each input image.

### 2.4 Morphometric feature extraction

Morphometric features were extracted from the filled vessel mask using the regionprops table function from the scikit-image library (skimage.measure). The label() function was applied to the filled mask to assign a unique integer identifier to each connected component. The following properties were computed for every labelled vessel:

#### Area

The total number of pixels enclosed within the vessel region.

#### Major axis length

The length of the longest axis of the ellipse best fitting the vessel region.

#### Minor axis length

The length of the shortest axis of the best-fitting ellipse.

#### Eccentricity

The ratio of the distance between the foci of the best-fitting ellipse to its major axis length; ranges from 0 (circle) to 1 (line segment).

#### Orientation

The angle in radians between the major axis of the best-fitting ellipse and the horizontal axis of the image.

#### Centroid

The (row, column) pixel coordinates of the geometric centre of the vessel region in image space, enabling spatial mapping of vessels across the tissue section.

### 2.5 QuPath extension and graphical interface

To make the Python pipeline accessible within routine digital pathology workflows, VeSpA was implemented as a QuPath extension developed in Java. Once installed, VeSpA is accessible from the QuPath Extensions menu under “Vessel Segmentation”, which opens a dedicated graphical user interface. The interface allows users to select the processing scope, including selected annotations, TMA cores, or the whole image, configure segmentation parameters, execute the Python backend, and import vessel objects directly into the QuPath hierarchy.

The graphical interface is organised into a sidebar and a central workspace (Figure 1). The sidebar provides real-time status indicators, including image availability, current selection or TMA-core availability depending on the selected processing mode, and Python environment readiness. A dedicated “Configure Python” button opens a configuration dialog where users can specify the Python executable, validate the environment, and check required dependencies. The central workspace contains the input region selection, segmentation presets, quick controls, run panel, live progress bar, log window, and advanced tuning options.

**Fig. 1:**
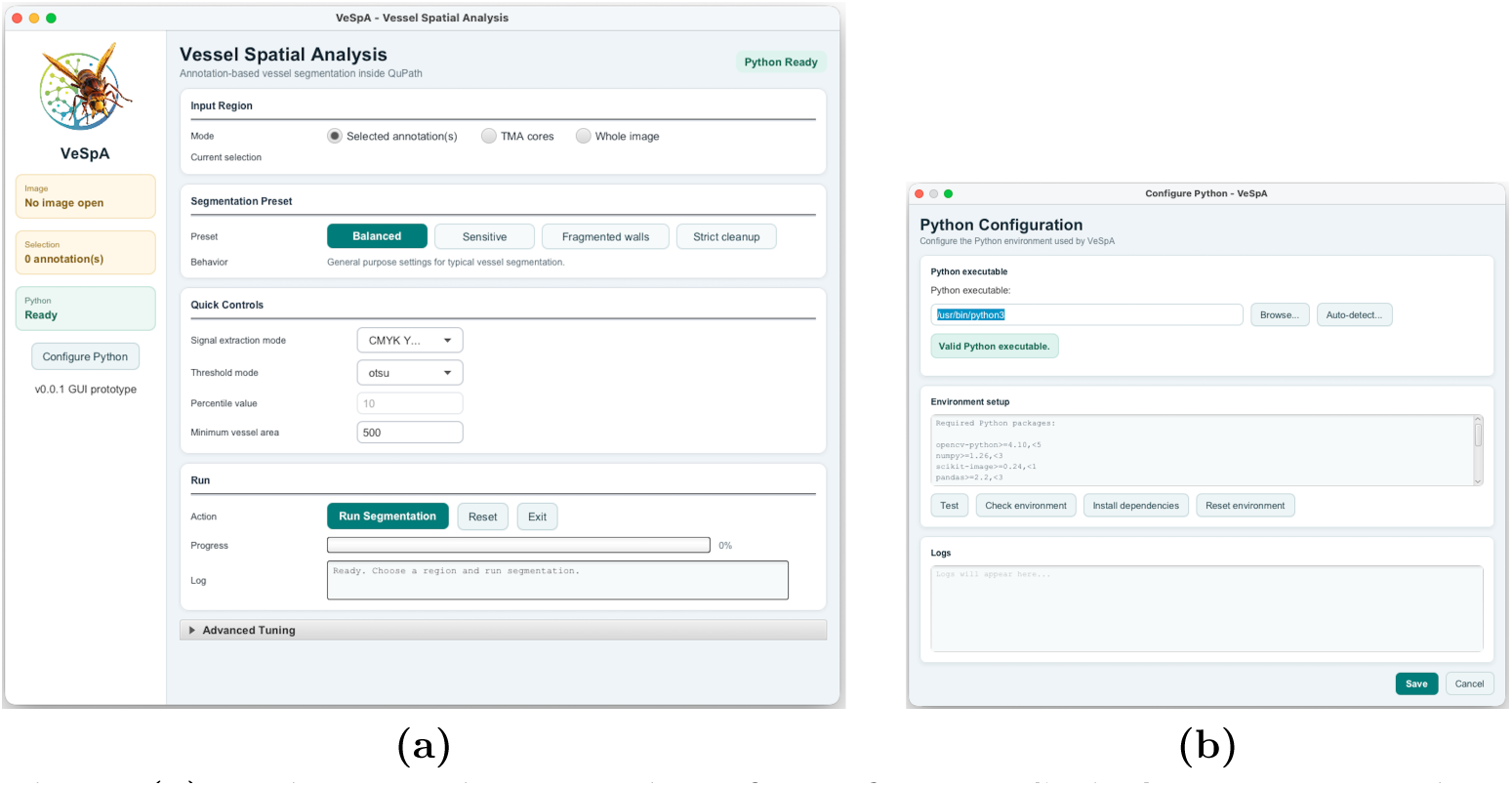
**(a) Main graphical user interface of the VeSpA QuPath extension.** The VeSpA GUI provides a sidebar with image, selection or TMA-core availability, and Python environment status, together with a central workspace for region selection (selected annotations, TMA cores, or whole image), segmentation presets, quick controls, live progress display, log output, and advanced tuning parameters. The updated interface also includes configurable signal extraction, allowing either CMYK Yellow preprocessing (default) or optional DAB stain deconvolution for H-DAB images. **(b) Python configuration dialog of the VeSpA QuPath extension.** Interface used to configure and validate the Python environment required for running the VeSpA pipeline. Users can specify or auto-detect the Python executable, test the interpreter, check required dependencies, install missing packages, and reset the environment when needed. The dialog ensures reliable communication between the QuPath extension and the Python backend, enabling stable execution of segmentation and data exchange within the workflow.

The *Input Region* panel allows users to select the processing scope, including selected annotations, TMA cores, or the entire image. The *Segmentation Preset* panel provides four predefined configurations: Balanced (default, general purpose), Sensitive (lower thresholds for smaller vessels), Fragmented walls (larger closing kernel for disrupted vessel walls), and Strict cleanup (higher area thresholds to reduce noise). These configurations automatically populate the parameter fields and allow rapid adaptation to different image characteristics. Frequently adjusted parameters are exposed in the *Quick Controls* panel, including signal extraction mode, thresholding mode, percentile value, and minimum vessel area. More detailed settings are available in the *Advanced Tuning* panel, including lumen detection, wall repair, morphological operations, and lumen expansion parameters.

The extension communicates with the Python backend by exporting selected image regions, passing validated parameters to the segmentation script, and reading the resulting vessel contours and measurements. In QuPath, the polygon contour coordinates of each vessel are used to reconstruct the vessel as a polygon region of interest classified as “Vessel”. Each “Vessel” object has the same morphometric measurements as in the Python code attached to the object’s measurement list, including area, major axis length, minor axis length, eccentricity, orientation, and centroid. When run on selected annotations, all vessel objects are added as child detections of the corresponding parent annotation, and summary statistics, including vessel count, mean and total area, mean axis lengths, mean eccentricity, and mean orientation, are appended to the parent annotation’s measurement list. The same parent-level summary logic is applied to valid TMA cores when TMA-core mode is used. In whole-image mode, vessel detections are imported directly into the image hierarchy without annotation or TMA-core parenting. An overview of the complete VeSpA segmentation pipeline is shown in Figure 2.

**Fig. 2:**
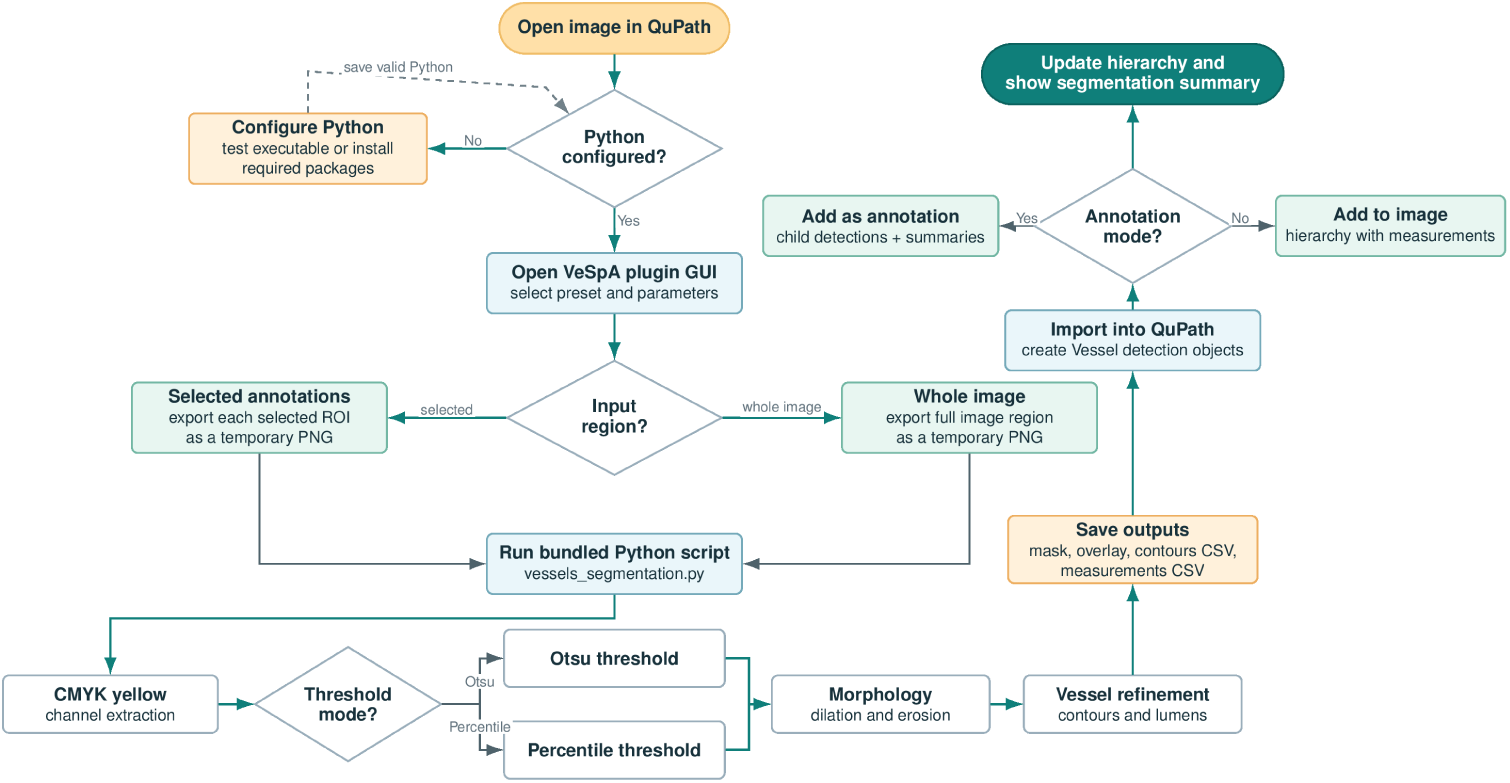
Overview of the Vessel Spatial Analysis (VeSpA) segmentation pipeline. The six sequential processing steps are shown with their key operations and configurable parameters.

### 2.6 Outputs

VeSpA generates two types of output depending on the interface used. In the standalone Python code, four output files are saved in a dedicated per-image subdirectory: the lumen-filled binary vessel mask, the green-channel visualisation overlay blended onto the original histological image, a morphometric measurements CSV file with one row per vessel and one column per descriptor, and a vessel contours.csv file containing polygon contour coordinates for downstream object reconstruction. The Python code also prints per-image summary statistics to the console, including total vessel count, mean area, total area, and mean eccentricity.

In the QuPath extension, selected annotations, valid TMA cores, or the whole image can be processed depending on the selected mode, generating vessel detection objects added to the QuPath hierarchy and summary statistics appended to the corresponding parent annotation or TMA core when applicable. These outputs make the results immediately accessible in QuPath’s measurement tables and scripts, while preserving compatibility with downstream statistical analysis.

### 2.7 Validation

Four images (ROIs) of 4000 *×* 4000 pixels containing at least 150 segmented vessels each were randomly selected from the available dataset. GG and SLR blindly and independently annotated all four sections, producing binary ground truth (GT) masks by manually delineating vessel regions. This yielded eight GT masks in total, two per section.

The GT masks were compared pixel-wise against the corresponding VeSpA segmentation mask with four different validation strategies. First, *per-rater comparison*: VeSpA masks were compared against each ground truth independently, yielding eight comparisons (four sections *×* two raters). For each comparison, a colour-coded difference image was generated encoding true positives in white, true negatives in black, false negatives (vessel present in GT but missed by VeSpA) in red, and false positives (vessel detected by VeSpA but absent in GT) in blue. Second, *inter-rater agreement* : the two ground truth masks for each section were compared against each other, yielding four comparisons. For the inter-rater difference images, neither mask was treated as reference; red pixels indicate regions annotated by GG only, and blue pixels indicate regions annotated by SLR only. Accuracy, Dice, and IoU are fully symmetric for this comparison; precision and recall are reported with GG as the nominal reference and should be interpreted as agreement rates rather than segmentation errors. Third, *intersection GT comparison*: for each section, a consensus ground truth was constructed as the pixel-wise intersection (logical AND) of the two raters’ masks, representing only regions on which both agreed to annotate a vessel; VeSpA was then compared against this conservative consensus. Fourth, *union GT comparison*: a permissive consensus ground truth was constructed as the pixel-wise union (logical OR) of the two raters’ masks, representing any region annotated; VeSpA was compared against this inclusive consensus. The intersection and union comparisons together bracket the range of possible performance relative to inter-rater disagreement. All four comparison strategies produced the following pixel-level metrics summarised as mean *±* standard deviation:

- Accuracy: (*TP* + *TN* )*/*(*TP* + *TN* + *FP* + *FN* )
- Precision: *TP/*(*TP* + *FP* )
- Recall (Sensitivity): *TP/*(*TP* + *FN* )
- Specificity: *TN/*(*TN* + *FP* )
- F1/Dice score: 2*TP/*(2*TP* + *FP* + *FN* )
- Intersection over Union (IoU): *TP/*(*TP* + *FP* + *FN* )

### 2.8 Benchmarking

Next, VeSpA was compared to two computer vision models through three further approaches on the same four sections: SAM (ViT-B checkpoint, automatic mask generation, prompt-based), YOLOv8-seg (pretrained yolov8n-seg.pt weights, zero-shot), and a hybrid VeSpA-SAM pipeline in which VeSpA-derived Y channel stain masks were used as spatial prompts for SAM boundary refinement. For the hybrid method, each image was first processed by VeSpA to generate binary candidate vessel regions; connected-component analysis was applied to these regions to derive one positive centroid prompt and one bounding box per component, which were then passed to the SAM predictor for boundary-guided refinement; final masks were combined by logical union and post-processed with morphological opening and closing. The same pixel-level metrics as in the previous validation were computed.

## 3 Results

### 3.1 Outputs

VeSpA successfully processed all selected annotations, generating individually labelled vessel objects for each detected structure. In representative CD31-stained sections, vessels were cleanly delineated as filled polygon regions, with lumens correctly identified and filled, albeit with difficulty in cases of particularly large lumen size or a very high degree of wall fragmentation. Small noise fragments arising from background staining or scanning artefacts were consistently excluded by the area filter, and the resulting binary masks contained no spurious detections at the default parameter settings. The contour overlay allowed rapid visual verification of segmentation quality, facilitating the identification of any segmentation errors at a glance. Representative step-by-step outputs illustrating each stage of the pipeline are shown in Figure 3.

**Fig. 3:**
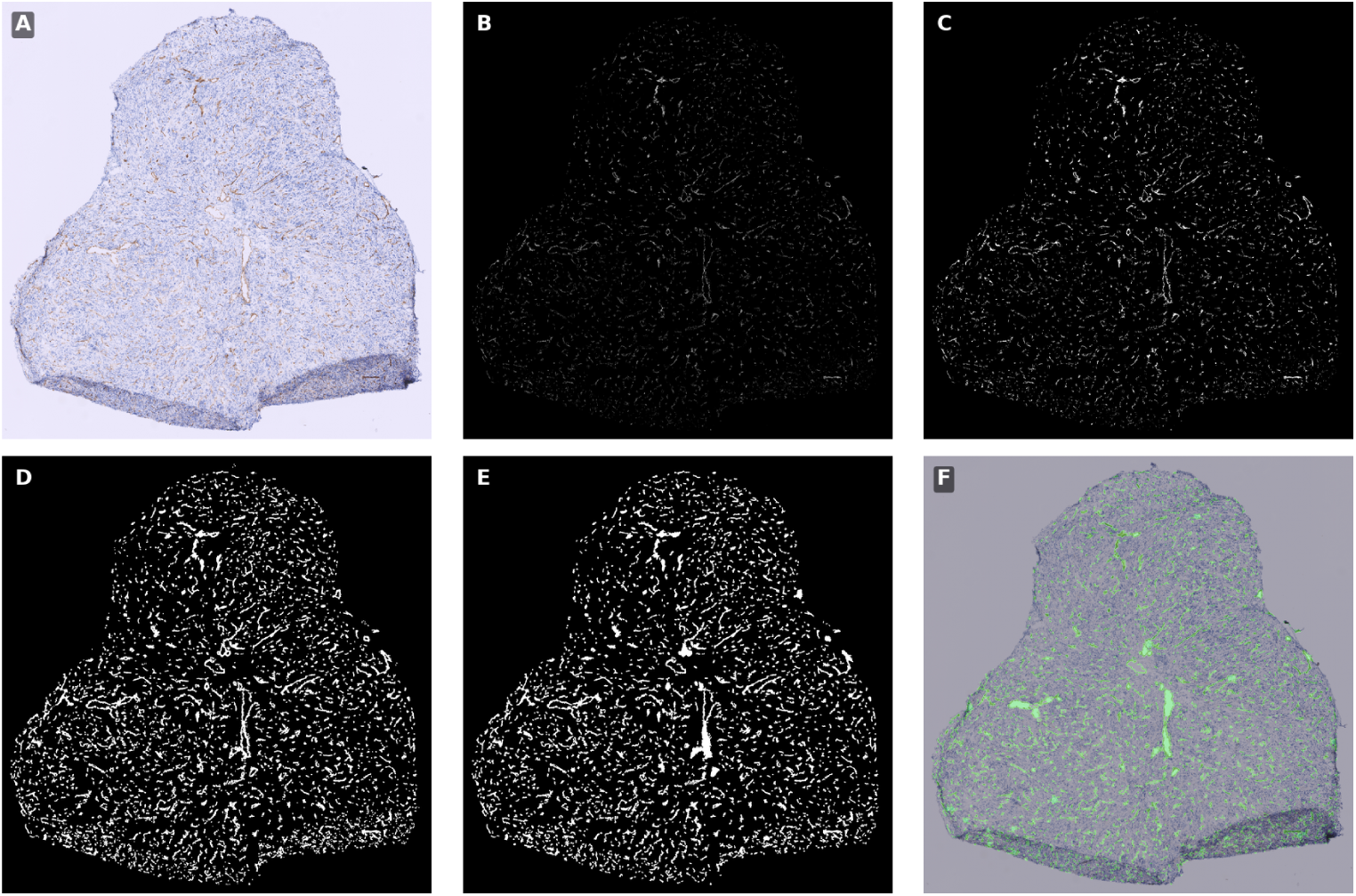
Representative step-by-step output of the VeSpA pipeline applied to a CD31-stained FFPE section. (A) Original image; (B) CMYK Yellow channel (default signal extraction mode); (C) binary mask after thresholding; (D) after morphological refinement; (E) after contour filtering; (F) final lumen-filled mask with overlay. An optional DAB stain deconvolution mode is also available in the current VeSpA implementation for H-DAB images.

### 3.2 Morphometric quantification

Across the dataset of 17 WSIs, VeSpA extracted 71,406 vessels in total and six morphometric descriptors for every detected vessel: area, major axis length, minor axis length, eccentricity, orientation, and centroid coordinates. The distributions of the five scalar descriptors and the spatial distribution of centroids across one example image are shown in Figure 4. Vessel area exhibited a right-skewed distribution consistent with the coexistence of small capillaries and larger venules in CD31-stained tumour tissue, with a median of 1,384 px^2^ (IQR 857–2,571). Major axis length showed a median of 60 px (IQR 43–94), while minor axis length showed a median of 32 px (IQR 26–42). Eccentricity values were broadly distributed between 0 and 1, with a median of 0.83 (IQR 0.72–0.91), reflecting the morphological heterogeneity of the tumoral vascular network, which includes both approximately circular cross-sections and elongated, tortuous structures. Orientation was approximately uniformly distributed across the full angular range (0°–180°), indicating the absence of a dominant vascular directionality in the analysed sections. The spatial scatter plot of centroid coordinates showed uniform vessel distribution across the ROI, with no systematic concentration at image borders, confirming that the segmentation did not introduce spatial biases. Processing time averaged 69–72 s per image on standard laboratory hardware (macOS 15.7.4, 1.1 GHz Intel Core i3 dual-core), confirming that VeSpA is computationally tractable for large dataset analysis without GPU requirements.

**Fig. 4:**
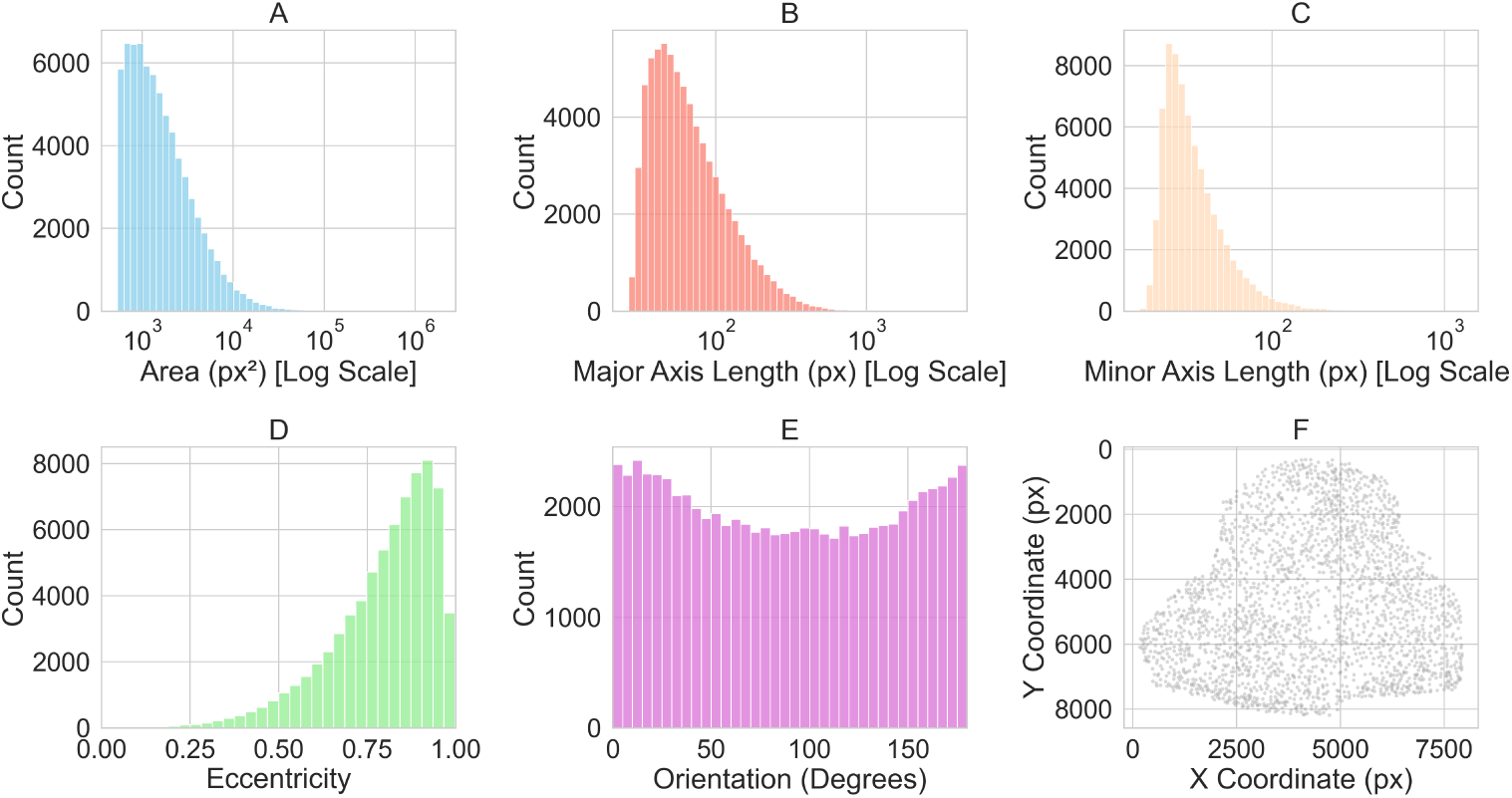
Distributions of morphometric descriptors extracted by VeSpA across all segmented vessels in the validation dataset. (A) Area; (B) major axis length; (C) minor axis length; (D) eccentricity; (E) orientation; (F) spatial distribution of vessel centroids across the tissue section.

### 3.3 Pipeline performance

Pixel-level comparison of VeSpA’s binary vessel masks against annotated ground truth masks was performed using four complementary strategies and yielded the results visible in Table 1. Inter-rater agreement figures treat GG as the nominal reference and reflect agreement rates rather than segmentation error. VeSpA’s per-rater Dice (0.743 *±* 0.040) was closely comparable to inter-rater Dice (0.768 *±* 0.019), indicating that the pipeline approaches the level of human agreement on this dataset. The intersection and union results bracket the performance range: VeSpA’s Dice of 0.743 is consistent across all four comparison strategies (range 0.742–0.768), and its recall is higher than the union benchmark (0.636) while its precision is lower (0.806 vs. 0.907), reflecting a balanced segmentation that closely tracks the region of inter-rater consensus. Difference maps for each validation strategy are shown in Figure 5. A summary comparison of VeSpA’s segmentation performance against ground truth across the four strategies is shown in Figure 6.

**Fig. 5:**
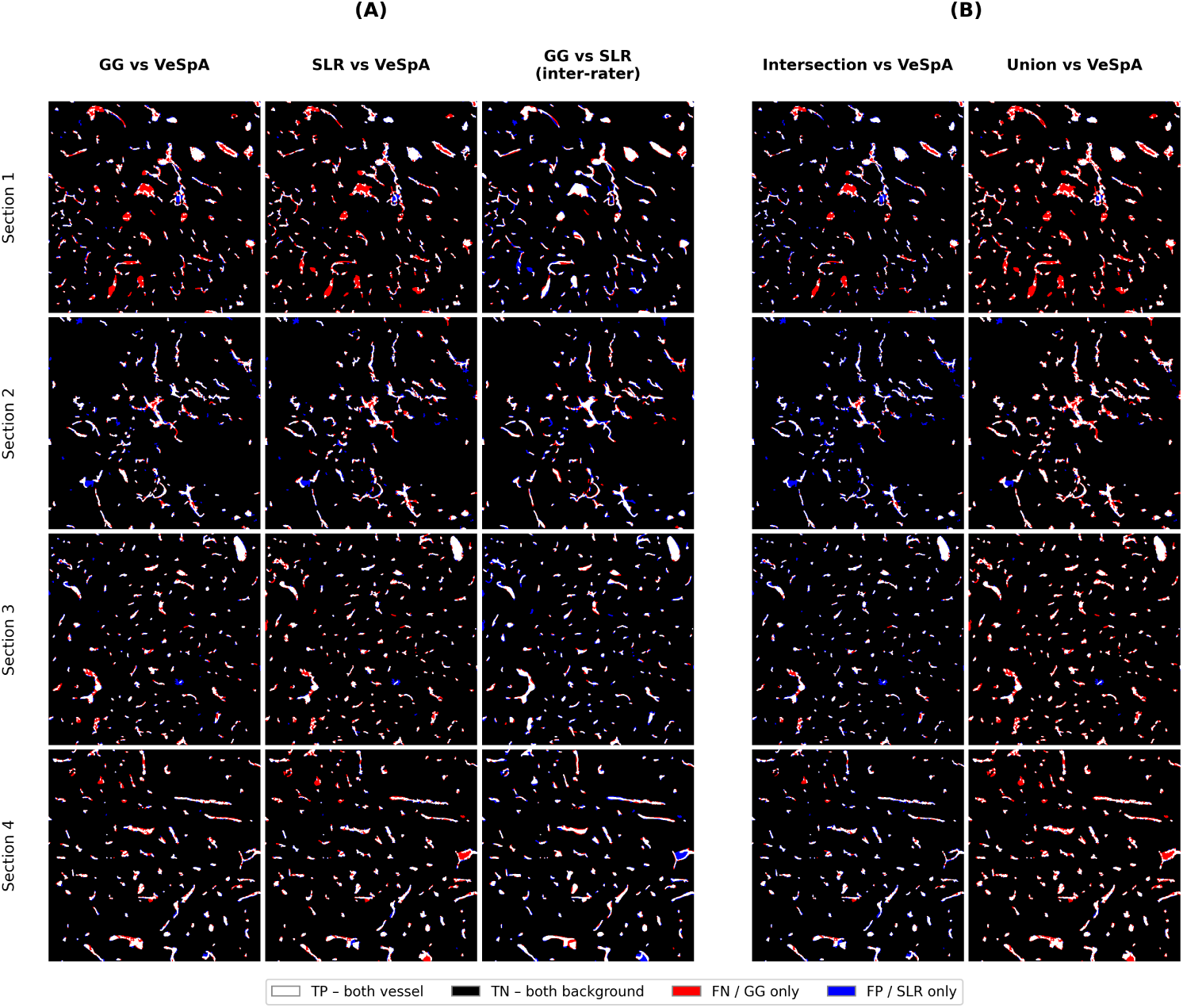
Pixel-level difference maps for each validation strategy across four tissue sections. Panel (A) 4 *×* 3 grid for per-rater and inter-rater comparisons; Panel (B) 4 *×* 2 grid for intersection and union consensus comparisons.

**Fig. 6:**
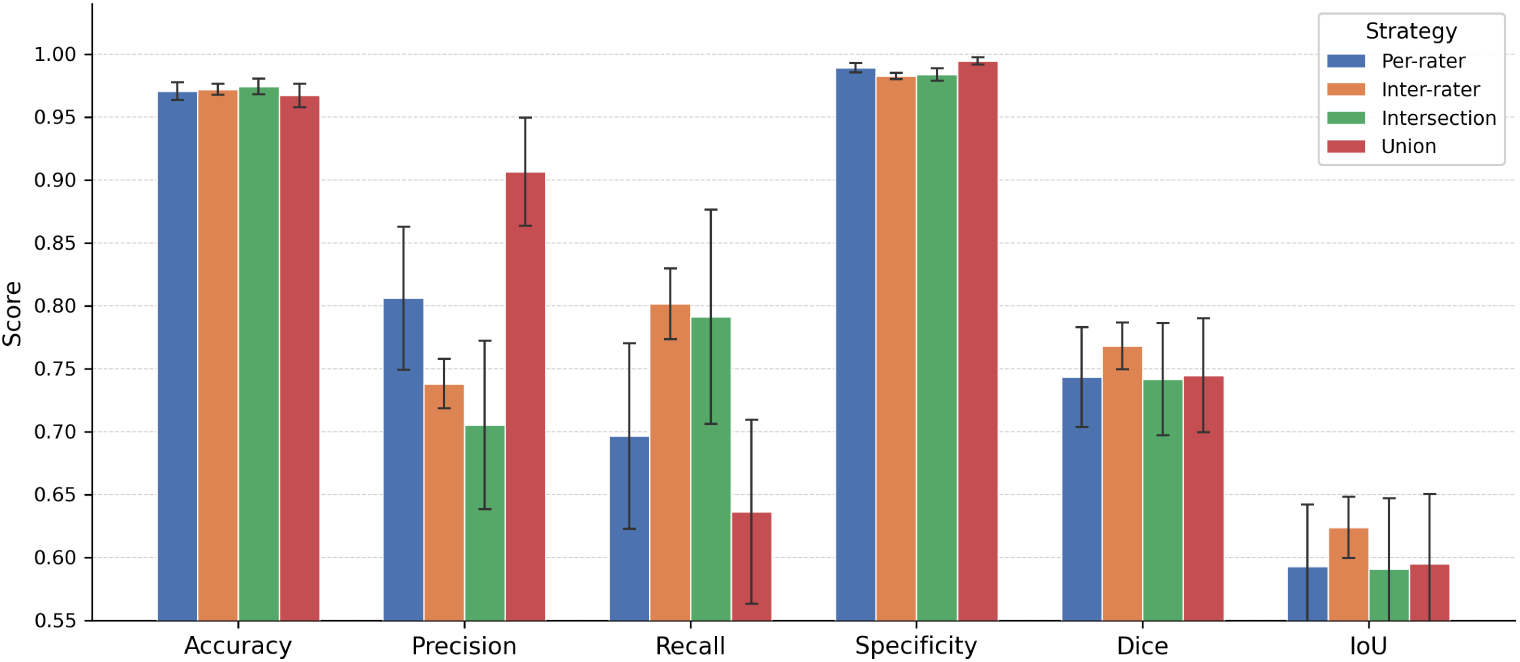
Comparison of VeSpA segmentation performance against pathologist ground truth. Bars show mean *±* SD across four tissue sections.

**Table 1:**
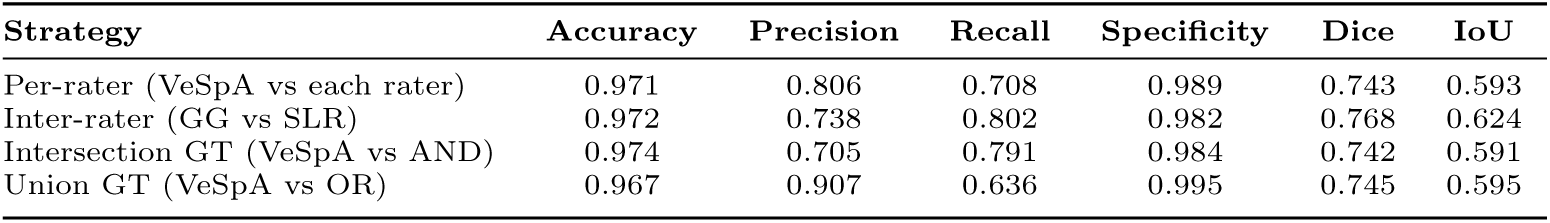
Validation metrics: VeSpA performance across per-rater, inter-rater, intersection, and union comparison strategies. Values are the mean across four tissue sections.

| Strategy | Accuracy | Precision | Recall | Specificity | Dice | IoU |
| --- | --- | --- | --- | --- | --- | --- |
| Per-rater (VeSpA vs each rater) | 0.971 | 0.806 | 0.708 | 0.989 | 0.743 | 0.593 |
| Inter-rater (GG vs SLR) | 0.972 | 0.738 | 0.802 | 0.982 | 0.768 | 0.624 |
| Intersection GT (VeSpA vs AND) | 0.974 | 0.705 | 0.791 | 0.984 | 0.742 | 0.591 |
| Union GT (VeSpA vs OR) | 0.967 | 0.907 | 0.636 | 0.995 | 0.745 | 0.595 |

### 3.4 Benchmarking

Benchmarking against SAM (ViT-B, automatic mask generation, prompt-based), YOLOv8-seg (pretrained yolov8n-seg.pt, zero-shot), and a hybrid VeSpA-SAM approach was performed on the same four sections. VeSpA achieved the highest mean Dice (0.743) and IoU (0.593), outperforming prompt-based SAM (Dice 0.299, IoU 0.179) and YOLOv8-seg (Dice 0.127, IoU 0.071) by a substantial margin. The hybrid VeSpA-SAM approach, in which VeSpA-derived stain masks were used as spatial prompts for SAM boundary refinement, ranked second overall with Dice 0.699, IoU 0.541, and the highest precision of the four methods (0.874); however, its recall (0.599) remained below VeSpA’s 0.708. Mean processing time per image was 2.58 s for VeSpA, 0.63 s for YOLOv8-seg, 25.23 s for SAM, and 43.46 s for the hybrid approach. Full metrics for all four methods are reported in Table 2 and Figure 7.

**Fig. 7:**
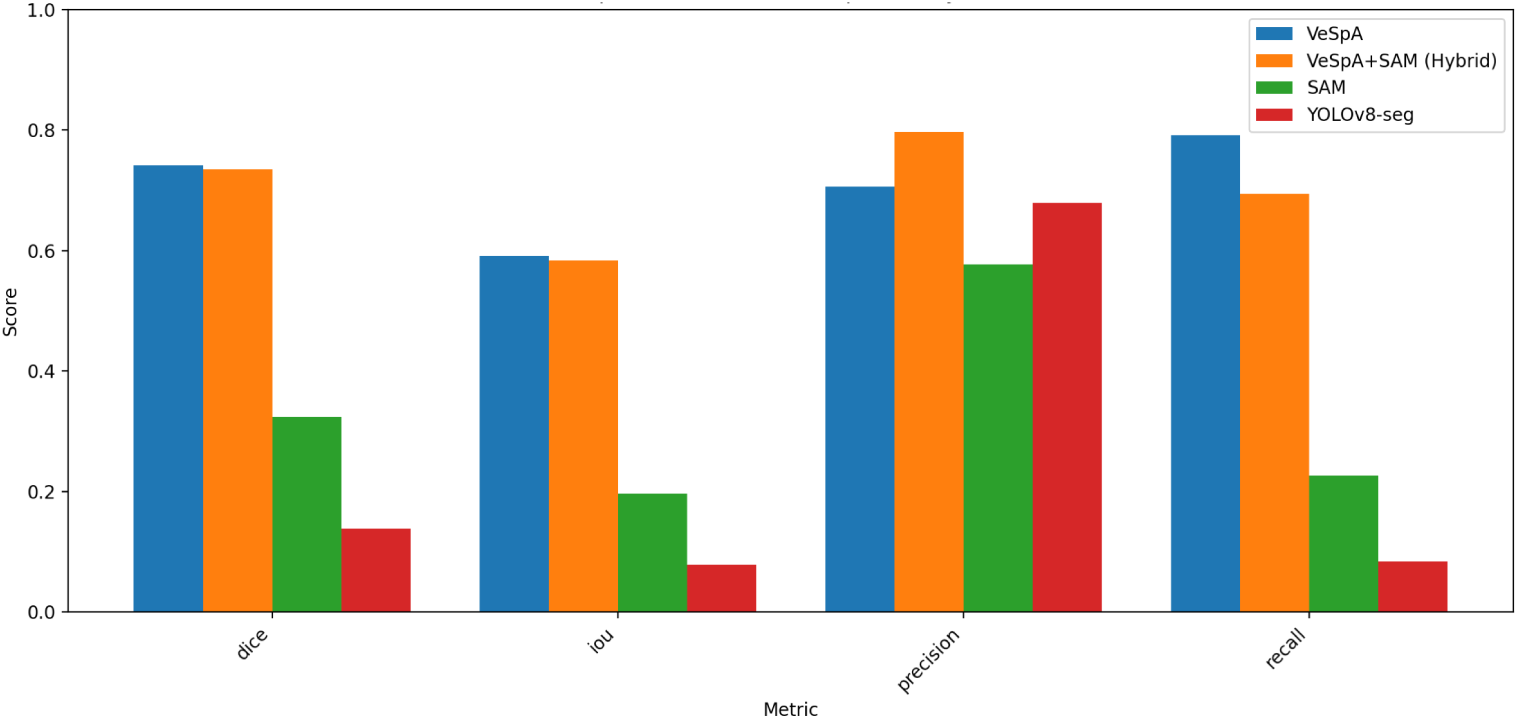
Benchmarking performance of VeSpA, VeSpA+SAM hybrid, SAM, and YOLOv8-seg across segmentation metrics.

**Table 2:**
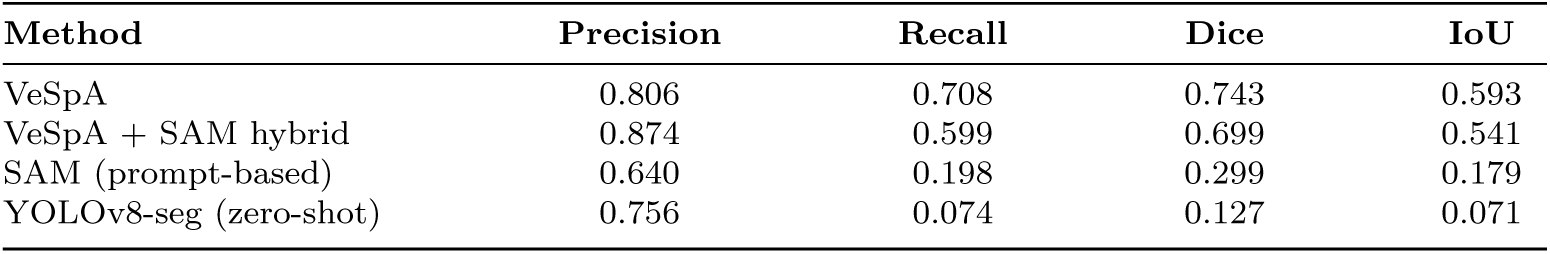
Benchmarking metrics: method comparison - VeSpA, VeSpA+SAM hybrid, SAM, and YOLOv8-seg. Values are the mean across four tissue sections.

## 4 Discussion

### 4.1 Principal findings and relevance

This study presents VeSpA as a QuPath-integrated, training-free workflow for automated vessel segmentation and morphometric quantification in CD31-stained WSIs. The main contribution of VeSpA is not only the generation of binary vessel masks, but the conversion of vascular staining into individual, measurable vessel objects that can be inspected, quantified, and analysed within the same digital pathology environment. This is important because vascular assessment in routine histology is often limited to qualitative review, manual counting, microvessel density estimation, or bulk stained-area measurements, whereas VeSpA enables reproducible single-vessel-level analysis across selected annotations, TMA cores, or whole-slide regions [12, 13, 29–34].

While platforms such as QuPath provide powerful tools for cellular quantification, annotation, and tissue analysis, dedicated validated workflows for vessel-level morphometric analysis in standard IHC-stained WSIs remain limited [12, 13, 15]. This gap is important because tissue organisation and disease progression depend not only on cellular composition, but also on the spatial relationship between cells, extracellular matrix, and vascular structures. By integrating vascular analysis into the QuPath environment, VeSpA allows vessel morphology and spatial distribution to be analysed alongside other digital pathology readouts, supporting a more complete characterisation of tissue architecture in oncological and inflammatory contexts [5–8]. The validation results indicate that VeSpA performs close to the level of inter-rater agreement between pathologists on the tested dataset. This is relevant because vascular annotation itself is subject to biological and interpretative variability, particularly in fragmented, tangentially sectioned, weakly stained, or irregularly shaped vessels [29, 35]. In this context, comparison with two independent pathologist annotations provides a more realistic benchmark than comparison with a single ground truth mask. The use of intersection and union consensus masks further shows that VeSpA behaves as a balanced segmentation approach, capturing the main region of pathologist agreement while avoiding excessive dependence on one annotator’s interpretation.

The benchmarking results also support the value of a domain-specific classical image processing strategy for this task. Although foundation models and generic segmentation networks are powerful tools, their yellow channel prompt-based and zero-shot application to CD31-stained histology did not match the performance of VeSpA in this dataset. This does not exclude the potential value of task-specific fine-tuning, as deep learning approaches have shown strong performance when trained or adapted for histological vessel analysis [23–27]. However, the present results show that a transparent and deterministic method tailored to the staining biology can remain highly effective, computationally lightweight, and easier to deploy in standard pathology research settings.

### 4.2 Relevance of the methodological design

The methodological design of VeSpA reflects a practical balance between interpretability, reproducibility, and usability. Rather than relying on a trained model, VeSpA uses stain-aware image processing steps that are transparent and directly inspectable. This is particularly relevant in computational pathology, where users often need to understand why a structure was segmented, adjust parameters for local staining conditions, and reproduce the same analysis across multiple slides or cohorts [12, 13, 28, 36].

The deterministic nature of the workflow is also important. Given the same input image and parameter settings, VeSpA produces the same segmentation output, which supports validation studies, collaborative analysis, and downstream statistical comparisons. This contrasts with workflows that depend heavily on model training, stochastic inference, or manual correction, where reproducibility may depend on training data, model version, hardware, or user intervention [23–27].

The integration of the workflow into QuPath further increases its practical relevance. Many image analysis methods remain difficult for pathologists or experimental researchers to use because they require command-line execution, data conversion, or separate scripting environments. By importing segmented vessels directly into the QuPath object hierarchy, VeSpA allows vascular measurements to be reviewed together with tissue annotations and other image-derived measurements. This makes vessel quantification more accessible and supports its use as part of broader digital pathology and spatial analysis workflows [12, 13].

The morphometric descriptors extracted by VeSpA should be interpreted in the context of 2D histological sectioning (Table 3). Among these measurements, minor axis length is particularly relevant because it provides a practical approximation of vessel caliber in cross-section and is less influenced than major-axis-dependent measurements by oblique or tangential cutting planes. For this reason, minor axis length may be especially useful when comparing vascular size distributions across annotations, tissue compartments, or experimental groups.

**Table 3:** Morphometric descriptors extracted by VeSpA for each segmented vessel.

| Descriptor | Definition | Biological interpretation |
| --- | --- | --- |
| Area | Total pixel area of the segmented vessel profile | Reflects the size of the segmented vascular profile; should be interpreted with awareness of sectioning angle, lumen filling, and vessel wall continuity. |
| Major axis length | Length of the longest axis of the fitted ellipse | Describes elongation of the 2D vessel profile; may be influenced by oblique or tangential sectioning. |
| Minor axis length | Length of the shortest axis of the fitted ellipse | Practical approximation of vessel caliber in 2D sections; relatively robust compared with major-axis-dependent measurements. |
| Eccentricity | Deviation from circularity of the fitted ellipse, ranging from 0 to 1 | Describes shape elongation or irregularity of the 2D profile; should not be interpreted alone as true 3D tortuosity. |
| Orientation | Angle of the major axis relative to the horizontal image axis | Describes image-plane orientation of the segmented profile; does not reconstruct the full 3D vessel trajectory. |
| Centroid | $(x, y)$ pixel coordinates of the geometric centre of the vessel region in image space | Enables spatial mapping of vessels and distance-based analyses within tissue compartments. |

Other descriptors remain informative but require careful biological interpretation. Vessel area reflects the size of the segmented vascular profile, but it may also be affected by sectioning angle, lumen filling, and vessel wall continuity. Major axis length and eccentricity describe the shape of the segmented 2D profile, yet elongated profiles may represent true vessel elongation, tangential sectioning, or partial vessel wall fragmentation. Similarly, orientation describes the direction of the segmented object within the image plane, but it should not be interpreted as a full measure of vascular trajectory in 3D tissue. VeSpA therefore provides these descriptors as quantitative features for comparative analysis, while biological interpretation should prioritise measurements that are most robust for 2D histology, particularly minor axis length, vessel count, area distribution, and spatial position.

The per-annotation output in QuPath and the three-file output in Python, binary mask, visualisation overlay, and morphometric CSV, are all explicitly designed to support both rapid quality control and bridge the gap between segmentation and downstream analysis. The structured CSV outputs are directly compatible with standard statistical environments, facilitating integration with spatial omics data modalities such as multiplex immunofluorescence, spatial transcriptomics, or imaging mass spectrometry. This structured output facilitates both visual quality control and systematic downstream analysis across large image datasets.

The validation framework employs four complementary comparison strategies, per-rater, inter-rater, intersection consensus, and union consensus, which together bracket the plausible range of VeSpA performance relative to human inter-rater disagreement and provide a more robust and informative assessment than single-rater comparison alone [35].

### 4.3 IHC markers for vascular visualisation

Immunohistochemical staining of endothelial cells with markers such as CD31 (platelet endothelial cell adhesion molecule-1, PECAM-1) or CD34 provides reliable and specific labelling, making these markers the gold standard for vascular visualisation in histological sections [37, 38]. While qualitative vessel evaluation on CD31-stained sections remains the reference approach in many research and clinical settings, it is inherently labour-intensive, subject to inter-observer variability, and impractical at the scale of WSIs containing thousands of vessels per slide [29]. The convenience of VeSpA lies in its automatic application to standard FFPE sections requiring only a standard endothelial DAB-stained section with no additional staining protocols [28, 39]. CD31 was selected as the primary endothelial marker because of its strong and specific labelling of endothelial cells lining vessel walls, providing reliable vessel–background contrast across a wide range of tissue types and fixation protocols. One important consideration is that CD31 also labels non-endothelial cells in certain contexts, including platelets, macrophages, and natural killer cells; users should therefore verify that the staining pattern in their tissue is predominantly endothelial before applying VeSpA. CD34 represents a broadly applicable alternative endothelial marker compatible with the same pipeline, although it can be less specific in certain contexts.

### 4.4 Comparison with existing tools

It is instructive to contextualise VeSpA within the landscape of existing vascular image analysis tools. A broad methodological perspective on this landscape is provided by Moccia et al. (2018), whose systematic review of vessel segmentation algorithms across modalities underscores that no single approach dominates across multiple imaging contexts [15]. A large body of work has addressed vascular quantification in fluorescence microscopy images, spanning both 2D and 3D contexts. Tools such as RAVE, AngioTool, AutoTube, and REAVER were developed for network-level analysis of fluorescently labelled vasculature, extracting parameters including vessel length, branching index, lacunarity, and segment connectivity [17–20]. SproutAngio and VESNA extend this to 3D fluorescence volumes, targeting *in vitro* models such as fibrin bead assays, vascular organoids, and hydrogel cultures [21, 22]. Morin et al. (2015) complemented these efforts by leveraging confocal microscopy to quantify the spatial organisation and interconnectivity of microvessels within synthetic scaffolds, while Adamo et al. (2022) proposed a deterministic algorithm based on spatial filtering and morphological feature extraction for the quantification of microvessels [16, 40]. While these tools have collectively advanced the field considerably, their dependence on fluorescence labelling and, in several cases, on 3D imaging or tissue clearing protocols makes them poorly suited to the analysis of standard FFPE material.

VeSpA is designed exclusively for 2D standard FFPE sections, which represent the most prevalent histological output in both research and routine clinical pathology. For this reason, IHC-based approaches are methodologically closer to VeSpA and merit more detailed consideration. Chantrain et al. (2003) were among the first to demonstrate computerised quantification of tissue vascularisation directly from high-resolution scans of whole tumour sections stained with chromogenic IHC, establishing that automated analysis of DAB-stained slides was both feasible and reproducible [30]. Sullivan et al. (2009) built on this by showing that automated microvessel area measurement from IHC-stained breast cancer sections is not only reproducible across observers but independently associated with patient prognosis, providing a strong clinical rationale for the kind of per-vessel area quantification that VeSpA performs [31]. Reyes-Aldasoro et al. (2011) developed a fully automatic algorithm for the segmentation and morphological analysis of microvessels in immunostained tumour sections, demonstrating that classical image processing without machine learning can reliably delineate vessel boundaries in IHC data and extract shape descriptors [32]. Kather et al. (2015) took a density-oriented approach, generating continuous spatial maps of microvessel density across WSIs to identify angiogenic hotspots, while Marien et al. (2016) systematically developed and validated a histological method for microvessel density measurement in WSIs of cancer tissue, addressing inter-observer variability and scanner dependence [33, 34]. Together, these studies establish that IHC-based vascular quantification in WSIs is analytically rigorous and clinically meaningful, but they also reveal a consistent gap: most existing IHC pipelines quantify vessel density or total stained area rather than providing per-vessel morphometric descriptors that reflect the shape and organisation of individual vessels.

Deep learning has more recently been applied to vessel segmentation in both histological and radiological images, with promising results. Yi et al. (2018) trained fully convolutional networks to predict microvessels directly from H&E-stained images without any IHC marker, demonstrating that vessel morphology encodes sufficient visual signal for learning-based detection [23]. Timakova et al. (2023) applied AI-assisted detection to blood vessels in WSIs in an oncological pathology setting, reporting gains in consistency and throughput over manual workflows [24]. Whitmarsh et al. (2025) presented perhaps the most directly comparable deep learning approach, applying semantic segmentation to CD31-stained breast cancer WSIs and quantifying multiple features of the tumour vasculature environment, achieving close agreement with expert annotation [25]. In the radiological domain, Praschl et al. (2023) demonstrated U-Net-based vessel segmentation in murine brain micro-MRI data with very limited training sets, highlighting both the versatility and the data dependency of supervised architectures [26]. DANEELpath integrates deep learning vessel segmentation within a QuPath-based spatial analysis framework, adding Voronoi influence zone mapping and centre-periphery analysis [27]. Collectively, these deep learning methods demonstrate high segmentation accuracy, but require significant computational resources and annotated training sets for fine-tuning that are difficult to compile for histological WSIs at scale. These are practical barriers that VeSpA’s training-free approach is specifically designed to avoid. VeSpA’s classical image processing approach achieves robust pixel-level segmentation performance on CD31-stained WSIs without any training data or GPU requirements, making it immediately deployable on standard laboratory hardware. Importantly, the deterministic nature of the segmentation strategy means that identical inputs will always produce identical outputs, a property that is not guaranteed by stochastic deep learning inference and that is of particular value in reproducible research contexts. Each step of the pipeline is transparent and directly interpretable: the effect of each operation on the image can be inspected at any stage, facilitating troubleshooting, parameter adjustment, and methodological reporting.

Comparison with SAM and YOLO-based segmentation contextualises VeSpA within the landscape of both classical and foundation model approaches. VeSpA substantially outperformed prompt-based SAM and YOLOv8-seg, and its per-rater Dice of 0.743 was closely comparable to the inter-rater Dice of 0.768, indicating that the pipeline approaches the level of human agreement. An important caveat is that neither SAM nor YOLOv8-seg was fine-tuned for histological vessel segmentation; the reported results therefore reflect zero-shot transfer performance rather than the upper bound of these model families, and task-specific fine-tuning may narrow the observed gap. The hybrid VeSpA-SAM pipeline demonstrates that integrating classical colour-space priors with SAM boundary refinement substantially reduces false positives relative to generic SAM, achieving the highest precision of the four methods; however, this comes at the cost of lower recall and a runtime of 43.46 s per image, nearly seventeen times slower than standalone VeSpA, making it impractical for large-scale batch processing in its current form. Collectively, these findings demonstrate that a training-free, domain-specific classical pipeline can substantially outperform zero-shot general-purpose models for this task while remaining computationally efficient on standard laboratory hardware without GPU requirements.

### 4.5 Limitations

The present study establishes VeSpA for its intended use case: vessel segmentation and morphometric quantification in standard CD31-DAB-stained FFPE WSIs. The current validation was performed on a defined set of oncological tissue specimens and one endothelial marker. This focused design allowed controlled assessment against independent pathologist annotations, but further testing across additional tissue types, scanner platforms, staining protocols, and endothelial markers will be useful to define the full range of applicability. The modular structure of VeSpA supports this extension because the colour extraction, thresholding, morphological refinement, and feature extraction steps can be independently adapted without changing the overall workflow. A second consideration is that VeSpA is currently optimised for brown DAB chromogenic IHC. This is appropriate for many routine CD31 and CD34 workflows, and the current implementation now supports both CMYK Yellow extraction as the default preprocessing mode and optional DAB stain deconvolution for H-DAB images. Nevertheless, alternative chromogens, endogenous pigments, or laboratory-specific staining characteristics may still require adjustment of the signal extraction step. Rather than representing a limitation of the general concept, this defines a clear route for further development: additional signal-extraction presets could be implemented for red chromogens, pigment-rich tissues, or site-specific staining protocols.

A third point concerns parameter selection. The default settings were selected to provide robust performance in the tested CD31-DAB images, but histological images can vary in staining intensity, vessel wall continuity, tissue background, and scanning resolution. For this reason, VeSpA exposes key parameters through the graphical interface and includes predefined presets for common segmentation scenarios. This design allows users to tune the analysis when necessary while keeping the workflow accessible to non-programmers.

Finally, VeSpA provides vessel-level morphometric measurements from 2D histological sections and does not aim to reconstruct full 3D vascular topology. Features such as branching architecture, vessel tortuosity, and longitudinal vessel course are better assessed using dedicated 3D or network-based imaging approaches. In contrast, the strength of VeSpA lies in reproducible quantification of vessel objects in routine 2D FFPE sections, where vessel count, minor axis length, area distribution, and spatial position can provide useful and scalable descriptors of vascular architecture [21, 22].

### 4.6 Future directions

Future developments of VeSpA will focus on further improvement of segmentation accuracy, particularly for vessels with low-contrast staining [26, 27].

Additional signal-extraction presets for slides stained with red chromogens (HRP red / AEC) or containing endogenous pigments such as melanin or haemosiderin represent a prioritised near-term development. Although the current implementation already supports optional DAB stain deconvolution, further extensions would involve adapting the signal extraction step using complementary colour-space or stain-separation strategies, and extending the lumen-filling and contour-filtering steps accordingly. Such an extension would substantially broaden VeSpA’s applicability to alternative IHC protocols and tissue types where brown DAB staining is not feasible. A further planned development is the extension of VeSpA from per-vessel morphometric analysis to vascular network analysis. In the current implementation, each segmented vessel is treated as an independent object; a network-level extension would detect physical connections between vessels, identify branching points and vessel endpoints, measure the length and tortuosity of individual vessel segments between junctions, and characterise global properties of the vascular tree such as branching density and network complexity. This would bring VeSpA closer to the topology-oriented metrics provided by tools such as VESNA and SproutAngio, while retaining the per-vessel morphometric output that is specific to VeSpA [21, 22].

The benchmarking results show that a hybrid approach combining VeSpA colour priors with SAM boundary refinement achieves higher precision than VeSpA alone, suggesting that prompt-guided foundation model integration is a promising direction; reducing the hybrid method’s current runtime of 43 s per image, for example by pruning prompts, merging nearby components before SAM refinement, or batching prompts, represents a tractable engineering goal. Supervised fine-tuning of YOLO segmentation on task-matched CD31 histological annotations represents an additional avenue, as the current benchmark used pretrained weights without domain adaptation. An additional planned development is the integration of spatial correlation analysis with data from other markers. The use of Voronoi-based influence zones in DANEELpath to correlate vessel distributions with marker expression in adjacent tissue compartments points toward the broader potential of spatially resolved vascular analysis, and future integration of analogous spatial tools with VeSpA represents a natural direction for development [27]. This would allow users to ask spatially resolved questions such as how a given marker, for example a proliferation marker, a hypoxia indicator, or an immune cell population, distributes as a function of distance from the nearest vessel, or whether specific vessel morphologies co-localise with particular microenvironmental features. Such analyses are of direct relevance to tumour microenvironment characterisation and could be implemented by co-registering the VeSpA vessel mask with additional marker channels or WSIs from serial sections.

## 5 Conclusions

VeSpA provides a QuPath-integrated, training-free workflow for automated vessel segmentation and morphometric quantification in DAB-stained WSIs, supporting annotation-based, TMA-core-based, and whole-image analysis within a familiar digital pathology environment. By converting vascular staining into individual measurable vessel objects, VeSpA moves vascular assessment beyond qualitative review, manual counting, and bulk stained-area estimation, enabling reproducible single-vessel-level analysis within a familiar digital pathology environment.

The tool is designed for routine FFPE histology, requires no model training or GPU infrastructure, and produces interpretable outputs that can be inspected directly in QuPath or exported for downstream spatial analysis. Its validation against independent pathologist annotations and benchmarking against prompt-based SAM and zero-shot YOLOv8-seg support the value of a deterministic, domain-specific approach for this task.

VeSpA therefore fills a practical methodological gap in computational pathology: accessible, reproducible, and scalable vessel-level quantification from standard endothelial IHC-stained WSIs. Future extensions will broaden marker and chromogen compatibility and expand the workflow toward spatial and network-level vascular analyses.

## Code and data availability

The Python implementation of VeSpA and QuPath extension is available at: https://github.com/ComputationalPathologyLab/VeSpA. Documentation is available at: https://spatialvesselanalysis.github.io/. The benchmarking scripts and related validation workflow are available at: https://github.com/ComputationalPathologyLab/VeSpAbenchmarking.

## Declarations

### Ethics approval and consent to participate

Not applicable.

### Consent for publication

Not applicable.

### Author contribution

Giulia Grion: coding, validation, manuscript writing, review and revision. Rashid Hussain: coding, manuscript writing, review and revision. Filippo Emanuele Colella: coding, review and revision. Kirollos Roufail: coding, review and revision. Silvia Uccella: review and revision. Roberta Frapolli: review and revision. Cristina Matteo: review and revision. Ö mer Mintemur: coding, supervision, review and revision. Francesca Pennati: supervision, review and revision. Salvatore Lorenzo Renne: coding, supervision, validation, manuscript writing, review and revision.

